# Assessing Long-Term Stored Tissues for Multi-Omics Data Quality and Proteogenomics Suitability

**DOI:** 10.1101/2024.03.13.583805

**Authors:** Kyu Jin Song, Minsuh Kim, Yong Jin Heo, Kyung-Cho Cho, Jae-Won Oh, Dae Ho Kim, Chanwoong Hwa, Yeji Do, Seunghyuk Choi, Hee Sang Hwang, Kwoneel Kim, Kyunggon Kim, Seungjin Na, Eunok Paek, Joon-Yong An, Se Jin Jang, Min-Sik Kim, Kwang Pyo Kim

**Author notes:** **Correspondence:** (M.S.K.), (K.P.K.).

## Abstract

As research into the complexities of cancer biology deepens, the integration of multi-omics analyses has emerged as a powerful approach to unravel the complex molecular basis of cancers. However, challenges related to sample availability, including size, collection procedures, and storage duration, hinder the broad application of this methodology. Despite these limitations, there is a growing interest in exploring the potential of archived samples to expand the scope of multi-omics research. Our study aims to investigate the impact of storage duration on the measurment in genomic, transcriptomic, and proteomic profiles of archived samples, demonstrating their viability for advancing our understanding of cancer biology. To comprehensively address these trends and limitations, we systematically examined archived samples collected over a decade, focusing on their transcriptomic, proteomic, and phosphoproteomic attributes. Analysis revealed intricate patterns and dynamic shifts, especially in long-term transcriptomic data, with observed declines in read counts related to protein coding and gene coverage. However, these changes did not compromise the fundamental gene expression landscape. Proteomic result also demonstrated that storage period did not significantly influence proteomic measurement. Comparisons of housekeeping gene (HKG) and housekeeping protein (HKP) expressions unveiled consistent transcriptomic levels across samples, while distinctive proteomic disparities between tumor and normal tissues. In conclusion, the challenges posed by limited sample availability in multi-omics studies can be partially overcome through the strategic integration of archived samples. While technical shifts were evident in certain aspects of transcriptomic data, core gene expression patterns remained robust, and the functionality of essential transcription factors (TFs) and kinases remained unaffected. These findings underscore the potential of archived samples as valuable resources for multi-omics research, providing a broader landscape for investigating cancer biology and paving the way for more comprehensive insights into this intricate field.

## Introduction

Lung cancer stands as a global leading cause of cancer-related mortality, characterized by pronounced phenotypic and genetic heterogeneity^1, 2^. Histologically, lung cancers are categorized into small cell lung cancer (SCLC) and non-small cell lung cancer (NSCLC).

NSCLC can be further divided into distinct pathologic subtypes, including adenocarcinoma (ADC), squamous cell carcinoma (SC), adenosquamous carcinoma, and large cell neuroendocrine carcinoma, each exhibiting unique growth patterns, molecular characteristics, and responses to treatment. Hence, a comprehensive understanding of the molecular underpinnings of each subtype is imperative^3^.

Genome-wide genetic analysis has ushered in molecular-targeted therapies, offering improved survival rates for lung cancer patients. Nonetheless, therapeutic challenges like acquired resistance frequently manifest within a relatively short time frame^4^. Thus, unmet medical needs persist in NSCLC therapy, emphasizing the importance of a holistic comprehension of its biological mechanisms. Identifying diagnostic and prognostic biomarkers, as well as therapeutic targets, is pivotal for advancing treatment strategies.

Recent advances in the integration^5^ of genomics, transcriptomics, and proteomics for cancer proteogenomics has been conducted to discover risk factors, biomarkers for diagnosis and drug response prediction, and novel therapeutic targets in lung cancers^6–8^. This integrative approach bridges genomic aberrations with proteomic features and clinical outcomes, offering deeper insights into cancer biology and therapeutic opportunities.

However, challenges remain, particularly in ensuring the availability of surgical tissues that meet statistical significance criteria for proteogenomic analysis. To address this, the inclusion of biobank samples has proven advantageous. However, standard criteria for tissue storage duration in biobanks have yet to be established. There have been recent reports in the field of lung cancer proteogenomics that have utilized stored tissue samples with sample storage periods of 3 years^7, 9^ and studies with a sample storage period of 7 years^9^. Additionally, Lehtio et al. described the use of samples with a storage period of up to 5 years and 10 years for discovery and validation cohorts, respectively^10^. It is known that the risk of specimen deterioration in the process of collecting and storing surgical specimens affects the results of proteogenomics analysis^11^. These changes could impact the differentiation between normal and tumor tissue or confound the identification of specific subtypes.

Analyzing aged samples in multi-omics research also introduces concerns regarding measurement accuracy, with data quality assessment and control assuming paramount importance. In RNA-seq analysis, assessing gene expression levels and identifying differential expression patterns may be affected by tissue storage duration, potentially impacting genomic origin, biotypes, and gene coverage profiles. In proteomic analysis, the detection of peptide-spectrum matches (PSMs) can be influenced, possibly skewing the representation of biological trends. Post-translational modifications (PTMs) within PSMs may exhibit variations, complicating the characterization of key regulatory processes. Proteome-wide disruptions in protein cleavage patterns may further lead to discrepancies in peptide identification and quantification. Finally, the count of unique peptides, critical for bottom-up protein identification methods, should be considered, as sample aging may influence dataset reliability.

In this study, samples with a storage period of 4-12 years were examined and the effect of the storage period on proteogenomics research was confirmed through targeted next-generation sequencing (NGS) using the NextSeq platform (Illumina, San Diego, CA, USA) with OncoPanel AMC version 4 (OP AMC v4) and bulk RNA-seq with HiSeq 2500 platform (Illumina), and LC-MS/MS-based proteome analysis. Integrated measurements of DNA, RNA, proteins, and PTMs (phosphorylation and acetylation) were performed to measure the effect of storage period on multi-omics data quality. Our findings affirm the feasibility of employing biobank tissues stored for at least 12 years as valuable resources for uncovering novel therapeutic and diagnostic biomarkers at the levels of DNA, RNA, and protein.

## Experimental procedures and Statistical Rationale

### Human specimens

Cancer tissues and their matched non-neoplastic tissues were obtained from surgically resected specimens as part of the biobanking process for cancer at Asan Bio-Resource Center (Seoul, Korea) with patients’ consent. The research protocol was approved by the Ethics Committee of Asan Medical Center (Seoul, Korea), and the entire experimental protocol was conducted in compliance with the relevant institutional guidelines. The diagnosis of each case was confirmed by pathologists at Asan Medical Center. The tissue samples in this study were frozen and stored in the vapor phase of liquid nitrogen for 4 - 12 years. For this study, each sample frozen in a cryotube was pulverized, distributed by 47 - 55 mg per tube and divided into at least three tubes per sample (**Supplementary Table 1A**).

### Genomic analysis

DNA was extracted from pulverized tumor tissues with pulverized matched normal tissues using the DNeasy Blood & Tissue Kit (Qiagen, Germany) according to the manufacturer’s protocol. To evaluate the somatic mutations in the 18 paired tumour tissues, targeted next-generation sequencing (NGS) was performed using the NextSeq platform (Illumina, San Diego, CA, USA) with OncoPanel AMC version 4 (OP AMC v4). OP AMC v4 captured 323 cancer-related genes including 225 genes for entire exons, 6 genes for partial introns and 99 hotspots^12^. Genomic DNA (200 ng) was fragmented to an average of 250 bp by sonication (Covaris, Woburn, MA, USA) followed by size selection with Agencourt AMPureXP beads. A DNA library was prepared using the SureSelect XT custom kit (Agilent Technology) after checking DNA quality. Sequenced reads were aligned to the human reference genome (National Center for Biotechnology Information build 38) using BWA (0.5.9) with default options^13^. MarkDuplicates function from the Picard package (Broad Institute, Cambridge, MA; http://broadinstitute.github.io/picard, last accessed on February 14, 2018) was used to remove PCR duplicates. The deduplicated reads were realigned at known indel positions using the GATK IndelRealigner tool (Van der Auwera GA & O’Connor BD. (2020). Genomics in the Cloud: Using Docker, GATK, and WDL in Terra (1st Edition). O’Reilly Media.). Next, the base qualities were recalibrated using the GATK BaseRecalibrator tool. Somatic single-nucleotide variants and short indels were detected with an unmatched normal using Mutect2 and the SomaticIndelocator tool in GATK. Common and germline variants from somatic variant candidates were filtered out using the common dbSNP build 141 (found in > 1% of samples), Exome Aggregation Consortium release 0.3.1 (https://gnomad.broadinstitute.org/), the Korean Reference Genome database (http://coda.nih.go.kr/coda/KRGDB/index.jsp), and an in-house panel of normal variants. Final somatic variants were annotated using Variant Effect Predictor version 79^14^ and converted to the maf file format using vcf2maf (GitHub; https://github.com/mskcc/vcf2maf, last accessed on February 14, 2018). False-positive variants were curated manually using the Integrative Genomics Viewer^15^.

### RNA sequencing

Total RNA was extracted was extracted from pulverized tissues using the RNeasy Mini Kit (Qiagen) and a cDNA library was constructed using the TruSeq RNA Access Library Prep Kit (Illumina), starting with 1 ug of total RNA. All cases passed the cDNA library quality assurance tests (minimum requirement > 5 nM). Finally, 100 nt paired-end sequencing was performed using the HiSeq 2500 platform (Illumina). BCL files were transformed into FASTQ files using bcl2fastq package. FASTQ files were mapped to the human reference genome (National Center for Biotechnology Information build 38) using STAR v2.7.3a^16^ with recommended parameters from the GDC mRNA quantification analysis pipeline (https://docs.gdc.cancer.gov/Data/Bioinformatics_Pipelines/Expression_mRNA_Pipeline/, last accessed on February 14, 2018). HTseq-count^17^ was used for transcript quantification. The quantified gene expression was normalised using a fixed upper quartile normalization.

### Protein extraction and tryptic digestion

Human NSCLC and NAT samples were individually cryo-pulverized using a Cryoprep device (CP02; Covaris, USA). Each tissue was put into a cryovial (43047; Covaris, USA) on dry ice, moved to a Covaris tissue bag (TT1; Covaris, USA), placed in liquid nitrogen, and pulverized at an impact level of 3 for 30 s. Every tissue which was already cryo-pulverized was moved into cryovial (43047; Covaris, USA) only 50 mg amount for proteome. Fifty milligrams (wet weight) of cryo-pulverized human NSCLC and NAT samples were homogenized in lysis buffer at a ratio of about 300 μL lysis buffer for every tissue. 10 mL lysis buffer is composed of 5 % SDS, 100mM TEAB (pH 7.55), 100 × Halt™ Protease Inhibitor Cocktail (78430; Thermo Fisher Scientific, USA) 100μL, 10 × PhosSTOP™ (4906845001; Roche, Swiss) 1 EA, 5mg/ml PUGNAc (A7229, Sigma-Aldrich, USA) 14.3 μL for final 20uM concentration. Tissue lysis was performed by sonication using a Branson SFX550 Sonifier with 1/2” Horn, 550 W, 20 kHz, 240 VAC at a setting of pulse mode, on time 5 sec and off time 3 sec, total on time 30 sec, 25% at ice. The lysate was centrifuged at 15,000 x g at 4 °C for 10 min, and the supernatant was transferred to a new micro tube. To measure protein concentration, we used the Pierce™ BCA Protein Assay Kit (23227; Thermo Fisher Scientific, USA). Each 500 µg of protein was digested individually using the Suspension-Trapping (S-Trap) filter (C02-mini-80; Protifi, USA) method [10]. Proteins from each sample were denatured with 5% sodium dodecyl sulfate (SDS) in 100mM trietyl ammonium bicarbonate (TEAB). Reduction and alkylation were conducted by 20 mM 1,4-dithiothreitol (DTT) (10708984001; Roche, Swiss), boiled at 95 °C for 10 min, and 40 mM iodoacetamide (IAA) (I16125-10 g; Sigma-Aldrich, USA) for 30 min at room temperature (RT) in the dark. To form colloidal protein particles, added 7 multiple volumes binding buffer (90% methanol and 100mM TEAB) and 12% phosphoric acid (695017-100 ML; Sigma-Aldrich, USA) to its final concentration of ∼1.2% and gently inverting. Formed colloidal protein solution was loaded onto the S-Trap filter and centrifuged at 4,000 x g for 30s, and the flow-through was reloaded again. The filter was washed more than 3 times with 400 μL of binding buffer. Pierce™ Trypsin/Lys-C Protease mix, MS-grade (A41007; Thermo Fisher Scientific, USA) was used to digest samples at a 1:25 enzyme-to-substrate ratio and incubated 6 hr at 37 °C. Digested peptides were eluted by centrifugation at 4,000 x g for 30 s with 80 μL elution buffers which were sequentially performed in three steps as follows. 50 mM triethylammonium bicarbonate (TEAB) (90114; Thermo Fisher Scientific, USA), 0.2% formic acid (56302-50 ML; Fluka, USA) in H2O, and 50% acetonitrile and 0.2% formic acid. Collect all eluted tryptic peptides and dried using a Speed-Vac (Hypercool, Labex, Republic of Korea) and kept at −80 °C, until next used.

### Construction of the common reference pool

Global proteomics, phosphoproteomics and acetylproteomics of 15 patients were conducted in two batches using TMT 16 plex isobaric labeling reagent. For comparative analysis across all samples in two batches, common reference (CR) sample was needed in each TMT batches. Common reference sample is consisted of all the samples analyzed in the TMT and reflects all samples characteristics. 15 tumors and paired normal adjacent tumor tissues protein were pooled into one and same amounts of proteins as the other samples was performed with same digestion and distributed two 16 plex experiments. In batch, 134 N channels were used for CR and rest of channels were configured to contain approximately half of tumors and half of NATs.

### TMT-16 pro labeling of peptides

Peptides were labeled with 16-plex Tandem Mass Tag (TMT) pro reagents (A44520; Thermo Fisher Scientific, USA) as follows: 2 tumors with 126, 127 N, and 2 NATs with 127 C, 128 N, 2 tumors with 128 C, 129 N, 4 NATs with 129 C, 130 N, 130 C, 131 N, 4 tumors with 131 C, 132 N, 132 C, 133 N, 1 NAT with 133 C, 1 CR with 134 N TMT reagents. Others batch of TMT channel composition was opposed to it. Each peptide was diluted with 50 μL 100 mM TEAB and was added in 20 μL TMT reagents which was diluted in acetonitrile at 100 μg/μl. Mixture was incubated for 1 h at RT with gently shaking. 5% hydroxylamine was added into samples and incubated for 30 min at RT. For check labeling performance, a few aliquots of each channel were pooled into one and desalted by using Pierce C18 spin column (89870; Thermo Fisher Scientific, USA) for LC-MS/MS. Cut-offline to check TMT labeling efficiency is defined as 95 % peptides which has TMT labeling. Next, all TMT-labeled peptides in the batch were pooled and 1 % formic acid in water was added to bring the final concentration of acetonitrile in the sample to 4 % or less (pH 2∼3). Pooled samples were directly desalted using Sep-Pak C18 3 cc Vac Cartridge, 200 mg Sorbent per Cartridge, 55-105 µm (WAT054945, Waters) and elution were concentrated by vacuum centrifugation.

### Peptide Mid-pH fractionation

Labeled peptides were loaded on an analytical column (XBridge Peptide BEH C18 Column; 300 Å, 5 µm, 4.6 × 250 mm, 186003625; Waters™, USA) and guard column (SecurityGuard^TM^ cartridge C18, 4 x 3.0 mm; Phenomenex) for fractionation. A gradient was generated using a shimadzu HPLC LC20A operated with mobile phase A (10 mM triethylammonium bicarbonate in water, pH 8.5); and mobile phase B (10 mM TEAB in 90 % [vol/vol] acetonitrile, pH 8.5). The gradient was as follows: 0-8 min, 5% B; 8-65 min, 40%; 65-69 min, 44% B; 69-74 min, 60% B; 74-88 min, 60% B; 88-90 min, 5% B; 90-120 min, 5% B. Flow rate of mixture mobile phase is set to 1.0 mL/min. The separated peptides were collected every 0.91 min into 96 tubes and pooled into 24 fractions non-consecutively.

### Phospho-peptide Enrichment using IMAC

95% of each 24 fractions were used for phosphoproteomics and non-consecutively concatenated into 12 fractions. For enrichment, using immobilized metal affinity chromatography (IMAC) as previously described [11]. In brief, Ni-NTA agarose bead slurry (QIAGEN, 30410) was fully resuspended and took 20 μL per fraction. Spin beads down and removed flow through and washed with 1 mL HPLC water by gentle inverting and centrifuging at 1,000 x g, 1 min, 3 times. Incubate beads in 1.2 mL 100 mM EDTA for 30 minutes with end-over-end turning (SB3, Stuart, UK). Repeat wash bead steps by 1 mL HPLC water, 3 times. Incubate beads in 1.2 mL of a 10 mM FeCl_3_ aqueous solution for 30 minutes with end-over-end turning. Repeat wash bead steps again 3 times in 1 mL HPLC water. Resuspend beads into an appropriate slurry with 1:1:1 (Acetonitrile: Methanol: 0.01% Acetic acid) solvent and aliquot 10 μL beads for enrichment per fraction. For each of the 12 fractions, peptides were reconstituted to 0.5 μg/μL in binding/wash buffer (80% MECN, 0.1% trifluoroacetic acid) and coupled with aliquoted beads for 30 min at RT with gently mixing. After coupling step, the beads were spun down at 1,000 x g for 1 min and took off supernatant for acetyl enrichment. Beads were diluted by 80% MECN, 0.1% trifluoroacetic acid for desalting. Stage tips were used for desalting enriched peptides from IMAC. Stage-tips were pre-activated and equilibrated by methanol; 50% MECN, 0.1% formic acid; 1% formic acid sequentially. Beads were loaded on stage-tip and washed by 80% MECN, 0.1% trifluoroacetic acid and 1% formic acid. Flow through were also kept for acetyl enrichment. Next, elution buffer (500 mM KHPO4, pH 7) were used 3 times and 1% formic acid was added into stage-tips. All process used centrifugation at 1,000 rpm. After wash, elute peptide sample of stage-tip with 50% MECN, 0.1% formic acid, twice at 1,400 rpm. The eluted phosphopeptides were dried immediately by vacuum centrifugation for LC-MS/MS analysis.

### Acetyl-peptide Enrichment using Kac-Ab

Acetylated lysine peptides were enriched using an PTMScan® Acetyl-Lysine Motif [Ac-K] Kit (Cell signaling technology, 13416). IMAC flow through were concatenated into 4 fractions non-consecutively and dried down using vacuum centrifugation. Peptides were reconstituted with 1.4 ml of 1x diluted IAP buffer from 10x IAP buffer (5.78 g MOPS-NaOH, 1.461 NaCl, 0.8 g Dibasic sodium phosphate, and 0.08 g Monobasic sodium phosphate, adjust pH with Acetic acid ∼ 1.4 to 1.6 mL). Before incubated peptides with antibody, antibody was pre-washed 4 times by IAP buffers and separated to 2 aliquots. Peptides were incubated for 3 hours at 4°C with end-over-end turning (SB3, Stuart, UK). Coupled beads were washed 4 times with cold PBS, 3 times with cold HPLC water followed by elution with 100 μL of 0.15% TFA, twice. Eluents were desalted using C18 stage tips, eluted with 50% MECN, 0.1% formic acid and dried down using vacuum centrifugation directly.

### LC-MS/MS analysis

#### Proteome analysis

For LC-MS analysis, Ultimate 3000 RSLC nano system (Dionex, Germany) was coupled to Q Exactive HF-X hybrid quadrupole-Orbitrap mass spectrometer (Thermo Fisher Scientific, Germany) and equipped with a trap column (Acclaim™ PepMap™ 100 C18 LC Column, C18, 75 um x 2 cm, 5 μm, Thermo Scientific., 164564) for clean-up followed by an analytical column (EASY-Spray™ LC Columns, C18, 75 um x 50 cm, 2 μm, Thermo Scientific Inc., ES803A). Mobile phase flow rate was 300 nL/min, comprised of solvent A (0.1% formic acid and 3% acetonitrile in water) and solvent B (0.1% formic acid in 90% acetonitrile). The 185-minute LC-MS/MS method was set as follows: 0 to 12 min, 2% B; 12 to 14 min, 2% B; 14 to 17 min, 4% B; 17 to 120 min, 16% B; 120 to 150 min, 25% B; 150 to 155 min, 85% B; 155 to 163 min, 85% B; 163 to 165 min, 2% B, 165 to 185 min, 2% B. The analytical columns were maintained at 60 °C. The electric potential of electrospray ionization was set to 1.8 kV, and the temperature of the capillary was also set to 250 °C. MS1 spectra were acquired with a resolution 120,000, an automatic gain control (AGC) target of 3e6 and a mass range from 350 to 1500 m/z. The data-dependent acquisition in positive mode cycle was set to trigger MS/MS on up to the top 15 most abundant precursors per cycle at an MS2 resolution of 45,000 with dynamic exclusion time, 30 sec. And an AGC target of 5e5, a maximum injection time of 86 msec, fixed first m/z of 110 m/z, an isolation window of 0.7 m/z, and a normalized collision energy of 34 were set for MS2 acquisition. Peak Intensity threshold were set to 1e4 and the charge state of unassigned, 1, >6 was filtered out.

#### Phospho proteome and acetylproteome analysis

For LC-MS analysis, Ultimate 3000 RSLC nano system (Thermo Fisher Scientific) was coupled to Exploris 480 orbitrap mass spectrometer (Thermo Fisher Scientific, Bremen, Germany) and equipped with a trap column (Acclaim™ PepMap™ 100 C18 LC Column, C18, 100 um x 2 cm, 5 μm, Thermo Scientific., 164564) for clean-up followed by an analytical column (EASY-Spray™ LC Columns, C18, 75 um x 15 cm, 2 μm, Thermo Scientific Inc., ES901). Mobile phase flow rate was 300 nL/min, comprised of solvent A (0.1% formic acid with DW) and solvent B (0.1% formic acid in acetonitrile). The 185-minute LC-MS/MS method was set as follows: 0 to 14 min, 2% B; 14 to 17 min, 4% B; 17 to 120 min, 16% B; 120 to 145 min, 25% B; 145 to 150 min, 85% B; 150 to 158 min, 85% B; 158 to 160 min, 4% B. The analytical columns were maintained at 40 °C. The electric potential of electrospray ionization was set to 2.1 kV, and the temperature of the capillary was also set to 275 °C. Data-dependent acquisition positive ion mode was set with almost identical with global proteome except cycle time of 3 sec, an AGC target of 7.5e4, a maximum injection time of 120 msec and an HCD collision energy 38% in MS2 level.

### Mass spectrometry data analysis

#### Database search and quantitation using TMT ratios

The MS/MS spectra were searched against a composite database of uniport homo sapiens reference (January 2021; 97,679 entries) using Sequest HT search engine. Trypsin missed cleavage count is set up to 2. In common with global and PTM proteomic data, carbamidomethylation, alkylation of disulfide bonds in cysteine (+57.021 Da), TMT 16-plex modification of lysine, and N-termination (+304.313 Da) for static modification and oxidation of methionine (+15.995 Da) for variable modification. Dynamic modification of phosphorylation of serine, threonine, and tyrosine (+79.980 Da), acetylation of lysine (+ 42.01057 Da) added parameters to each PTM proteomic data. FDR was strictly evaluated as 0.01. In proteome discoverer 2.4 (Thermo Fisher Scientific, USA), reporter quantification was processed by each reporter ion abundance based on intensity. Normalization method with total peptide amount and scaling with on all average was used. Results on each proteome data, all identification protein and peptides were filtered with high confidence parameter for highly confident comparative quantification studies.

#### Open search for chemical modification discovery

Open search was conducted using MODplus v1.02^18^ to identify chemical modifications present in the samples. The unidentified MS2 spectra from the initial database search were searched, and their precursor m/z values were recalibrated by Proteome Discoverer during the initial search. The search parameters were set to 10 ppm for precursor mass tolerance, 0.01 Da for fragment mass tolerance, semi-tryptic enzyme specificity allowing for up to 2 missed cleavages, -1 to +2 for 13C isotope error, carbamidomethylation on cysteine residues, and TMT modification on the peptide N terminus and lysine residues for static modifications. All modifications in the Unimod database (v2020.10), excluding all amino acid substitutions, were considered for dynamic modification, and multiple modifications per peptide were permitted within the modified mass range of -150 to 350 Da. All PSMs were validated at 1% FDR using the MODplus FDR toolkit.

#### Two-component normalization of TMT ratios

To reliably perform quantitative comparison between samples, proteins or peptides that do not change are set to 0 in log2 TMT ratio distribution. Then we can calculate accurate regulation of conditions. A normalization corrects the data shape with this misaligned distribution to the zero-focus caused by systemic MS variation. 2-component Gaussian mixture model-based normalization algorithm was used to achieve this effect [6]. The two Gaussians N(μi1,δi1) and N(μi2,δi2) for a sample i were fitted and used in the normalization process as follows: the mode mi of the log-ratio distribution was determined for each sample using kernel density estimation with a Gaussian kernel and Shafer-Jones bandwidth. A two-component Gaussian mixture model was then fit with the mean of both Gaussians constrained to be mi, i.e., μi1=μi2=mi. The Gaussian with the smaller estimated standard deviation σi = min σi1, σi2 was assumed to represent the unregulated component of proteins/phosphosites/acetylsites and was used to normalize the sample. The sample was standardized using (mi), by subtracting the mean mi from each protein/phosphosite/acetylsite and dividing by the standard deviation δi.

### Dataset filtering and statistical analsysis

#### Identification of differentially expressed genes (DEGs)

To identify DEGs between conditions, we used ”DESeq“ function provided by DESeq2 R packages. Quantified transcript-level estimates from Salmon were imported to R using ”tximport“ and were converted to gene-level expression matrix. With this gene-level expression matrix, DESeqDataSet was generated using ”DESeqDataSetFromTximport“ function. Then, differential expression analysis based on the negative binomial distribution was performed using ”DESeq“ function. Of note, DEG analysis was conducted based on the formula below.

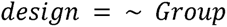

In this model, *Group* represents conditions (Tumor versus NAT) we want to compare.

#### Identification of differentially expressed proteins, phosphorylation, and acetylation

We identified differentially expressed proteins, phosphorylation, and acetylation using t-test analysis. After two-component normalization, we imputed missing values on data with k-nearest neighbors imputation method and used value of 5 for knn. These data were subjected to t-test analysis. Adjusted p-value was calculated using Benjamini–Hochberg method and adjusted p-value less than 0.05 was used as a criterion for differentially expressed proteins, phosphorylation, acetylation.

#### Gene set enrichment analysis

To obtain characteristic analysis information corresponding to tumor samples, we performed single sample Gene Set Enrichment Analysis (ssGSEA) (Barbie et al., 2009) for protein data. We computed normalized enrichment scores (NES) of gene sets. We used the implementation which contains ssGSEA available on GitHub (https://github.com/broadinstitue/ssGSEA2.0) using the command interface R-script (ssgsea-cli.R) using the following parameters:

➓ database: “h.all.v7.5.1.symbols.gmt”

➓ sample. norm. Type: “rank”

➓ output. score. Type = “NES”

➓ nperm=1000

#### TF activity estimation based on transcriptomic data

We also estimated TF activity from transcriptomics data. We used a curated collection of TF-target interaction provided by DoRothEA R packages. From DoRothEA gene regulatory network, “CollecTRI” were used to calculate TF activity scores. To identify tumor-specific TF activity, we used statistic value obtained from the DEG analysis (Tumor versus NAT) as an input. Also, we set minimum size of regulons to 10. We used ‘run_ulm’ function in decoupleR R packages to calculate TF activities.

#### Kinase activity estimation based on phosphoproteomic data

To understand the Subtype-specific kinase activities, we used decoupleR R package to infer kinase activity from the results of differentially expressed phosphorylation (DEPP) using prior knowledge network, such as databases for kinase/phosphatase and substrate interactions. We used prior knowledge network provided by PHONEMeS. Also, we set minimum size of regulons to 5, and provided 1,000 deregulated phosphorylation (each 500 phosphorylation for up-, down-regulated). There are a variety of methods to infer kinase activities and we used multivariate linear model (MLM)-based method as below:

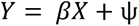

where the dependent variable *Y* represents the t-statistic measurements of phosphorylation from the results of the differentially expressed phosphorylation analysis, and the independent variable X is the connectivity matrix representing the associations with kinases. *X_ij_* equals 1 when phosphosite *i* is a known substrate of the kinase *j*, 0 otherwise. Ψ represents the normally distributed error of the fit and β represent the scores of the kinase activity. In decoupleR packages in R, we used ”run_mlm“ function to estimate kinase activity. P-value was obtained from multivariate linear model and adjusted p-value was calculated using the Benjamini–Hochberg adjustment.

## Results

### Multi-Omics Approaches for Tissue Sample

The analysis was conducted using a cohort of 18 patients who had been diagnosed with either lung cancer (89%, n=16) or non-lung cancer (11%, n=2) at the Asan Medical Center in Seoul, Korea (**Supplementary Table 1A**). Surgical procedures performed between 2009 and 2017 were utilized to collect both tumor and matched normal tissues (**Figure 1A**). Based on histological assessments, the patient population was categorized into four distinct groups: lung cancer-NSCLC-nonADC (’LNNA’, n=9), lung cancer-NSCLC-ADC (’LNA’, n=4), lung cancer-SCLC (’LSC’, n=3), and non-lung cancer (’NLC’, n=2) (**Figure 1A**). Within the LNNA subgroup, there were 5 cases of squamous cell carcinoma, 3 cases of adenosquamous carcinoma, and 1 case of pseudosarcomatous carcinoma. The LNA subgroup included 3 cases of solid carcinoma and 1 case of acinar cell carcinoma. The LSC subgroup consisted of 1 combined small cell carcinoma, 1 neuroendocrine carcinoma, and 1 small cell carcinoma. The NLC subgroup encompassed 1 case of thymoma, type B1, and 1 case of type B3 malignant thymoma. The TNM-based staging of the patients ranged from IB to IIIB, with approximately 30% of the cohort presenting with late-stage disease (IIIA: n=3, IIIB: n=2; see **Figure 1A**).

**Figure 1.**
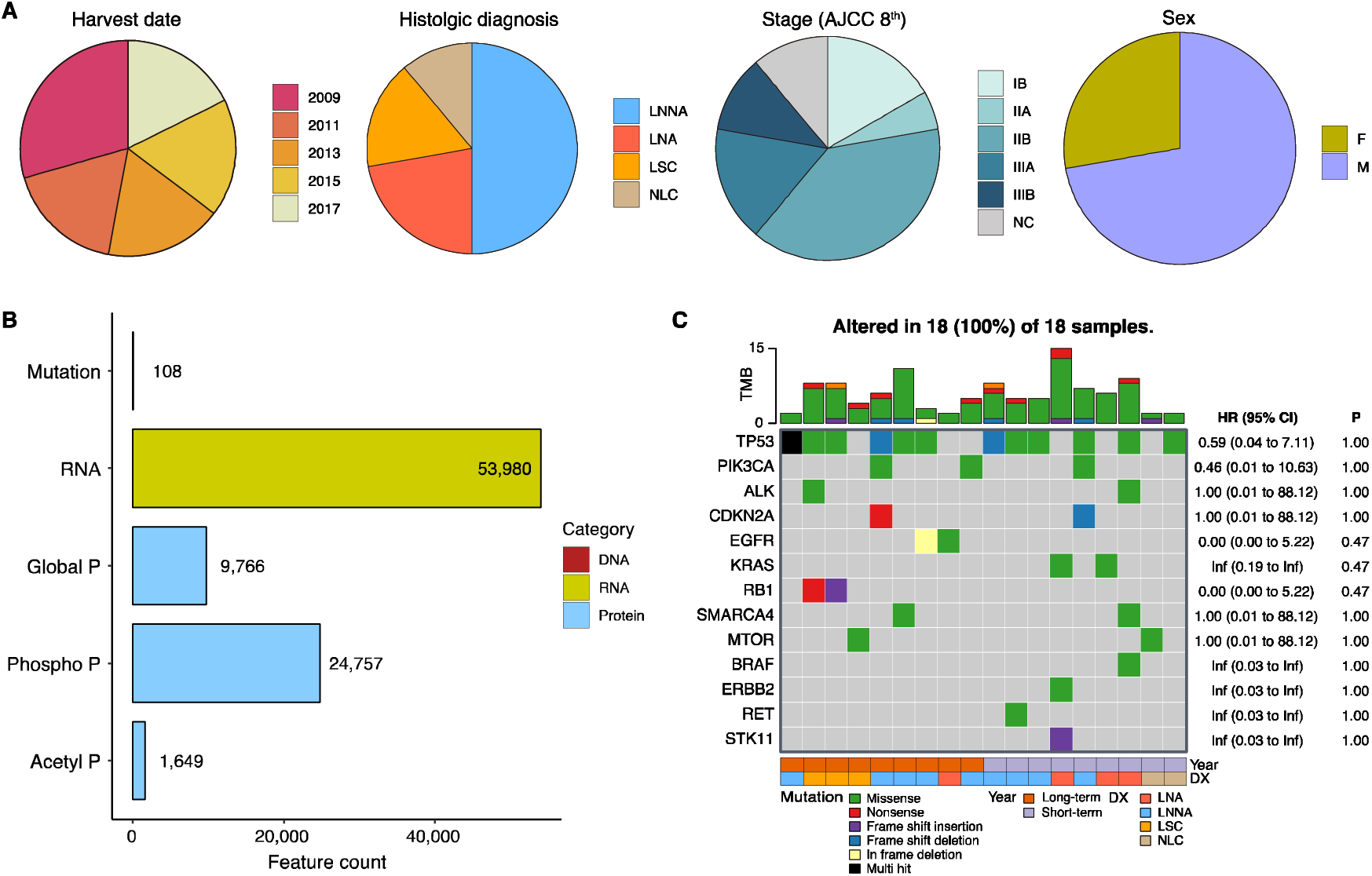
Distribution of samples and identification status in multi-omics study. **A.** Distribution of samples is illustrated based on the date of harvest, histologic diagnosis, tumor stage (AJCC 8^th^), and sex. **B.** Bar plot representing the feature identification status of samples which contains somatic small mutations, transcripts, proteins, phosphorylation sites, and acetylation sites. The color of each bar corresponds to the specific omics type. **C.** Oncoplot displaying results for each sample, indicating several known mutations (TP53, EGFR, KRAS, RB1, BRAF, and ERBB2…etc) associated with lung cancer. The bar plot at the top depicts the tumor mutational burden (TMB) for each sample, along with the sample frequency for each mutation. Table at the right panel shows fisher’s exact test result to mutation occurrence between long-term and short-term tumors.

We generated genomic, transcriptomic, proteomic, phosphoproteomic, and acetylproteomic datasets from the samples (**Figure 1B**). Each sample stored for long-term was pulverized and divided for multiple analyses. We used the samples that were divided by 50 mg to extract DNA, RNA, and protein. A genomic dataset was generated by targeted NGS covering 323 cancer-related genes in 18 tumor tissues and matched normal tissues. A total of 108 somatic mutations (SNV and indels) were observed in 18 tumors (**Figure 1B)** For transcriptomic analysis, we performed bulk RNA-seq for 17 tumors and their matched normal adjacent tissues (NATs) for deep coverage (80 - 100 M reads per sample), enabling gene expression quantification. We acquired 53,980 transcripts which cover 19,897 protein coding genes (**Figure 1B**). Using tandem mass tag (TMT)-based isobaric labeling, proteomic data were collected from 15 tumors and matched NATs. A total of 9,766 proteins, 24,757 phosphorylation sites, and 1,649 acetylation sites were observed for quantifiable features as a log_2_ ratio to the common reference (CR) sample (**Figure 1B**).

The common mutations in cancer tissues were successfully identified through targeted NGS analysis (**Figure 1C, Supplementary Table 1B**). These findings remained consistent when analyzing long-term samples, aligning with previous proteogenomic studies^7, 9, 10, 19^. The occurrence of TP53 mutations, a representative tumor suppressor, does not appear to be related to sample storage duration (p = 1, Fisher’s exact test). Similarly, in the case of EGFR mutations, there is no significant association with storage duration (p = 4.7 x 10^-1^, Fisher’s exact test).

### Assessment of RNA sequencing quality

In RNA-Seq analysis, read counts originating from exon regions are typically higher than those from intron regions. This disparity reflects the predominantly exonic nature of mRNA, as introns are generally spliced out during RNA processing. Nevertheless, the examination of intron regions is valuable for understanding transcriptional regulation and alternative splicing^20, 21^. Conversely, intergenic regions demonstrate the lowest read counts due to their lesser propensity for active transcription, often being deemed non-functional for protein-coding purposes. Across our samples, consistent patterns emerged, with minor variations between tumor and matched NATs (**Figure 2A, Supplementary Table 2A**). Notably, exonic alignment in NATs tended to be lower compared to intronic regions, primarily falling below 50% in total. These observations extend to various biotypes, including protein coding, lncRNA, miRNA, misc RNA, mt rRNA, pseudogene, and rRNA (**Figure 2B, Supplementary Table 2B**). Although overall gene coverage is relatively sustained despite extended storage, our results suggest a potential impact on gene coverage profiles indicating diminished transcript abundance (**Figure 2C, Supplementary Table 2C**). Notably, these effects, which are commonly used for expression analysis, were predominantly observed in NATs. Despite these changes in gene coverage profiles over the storage period, they do not significantly contribute to sample clustering. Principal Component Analysis (PCA) analysis was performed to evaluate the influence of storage duration, revealing unique clustering patterns primarily associated with the tissue sample type and pathology, rather than the storage period. (**Figure 2D**). Tumor samples form clusters in alignment with their pathology, displaying diverse distribution, while NAT samples exhibit relatively uniform gene expression values. Notably, two NATs from NLC cases appear to cluster with tumors, yet they remain distinct from their paired tumors, reflecting potential differences in tumor types. PC1 accounts for 22.6% of variance and predominantly segregates NATs from their paired tumor counterparts. Further analysis of PCA identified relevant variables contributing to expression clustering. Tissue type (tumor or NAT) emerges as the most pertinent factor (Bonferroni, adjusted p = 1.1 x 10^-4^), followed by pathologic diagnosis (Bonferroni, adjusted p = 5.5 x 10^-3^). Interestingly, the impact of sample storage duration, indicated by the sampling year, was not found to be statistically significant. Taken together, these findings indicate that samples stored for over a year suffice for gene expression analysis. Nonetheless, extended storage of NATs should be approached with careful consideration for potential future applications.

**Figure 2.**
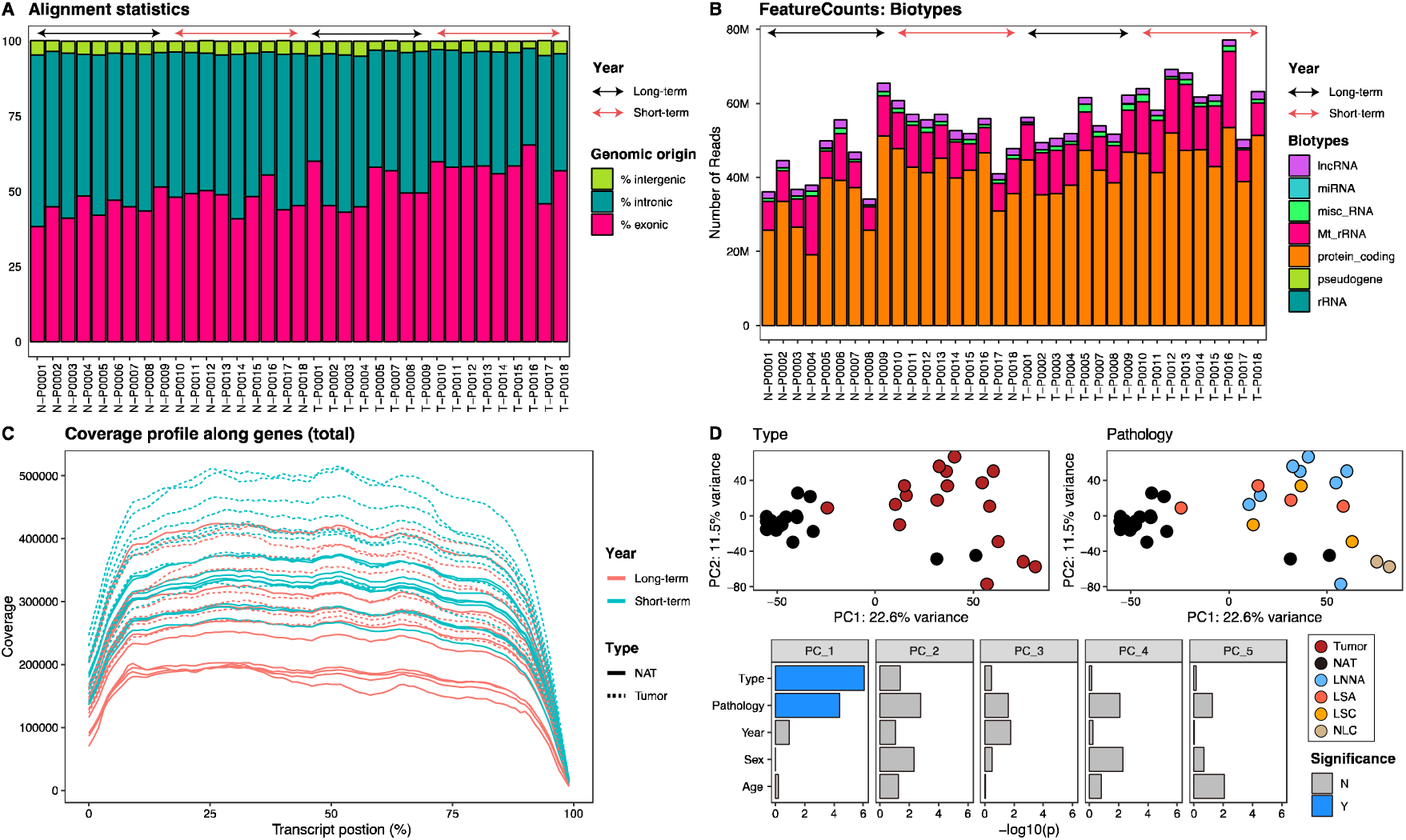
Assessment of read profile and gene expression using RNA-seq. **A.** Bar charts illustrating alignment statistics based on genomic origin of read counts in RNA-seq data. The color of arrows indicates the storage year. The y-axis represents the percentage of genomic origin in each sample. **B.** Bar chart depicting the distribution of biotypes in feature counts among samples. The color of bars indicates different biotypes, while arrows denote the year of storage. **C.** Line plot showing the mean distribution of coverage depth across the length of all mapped transcripts. The type of line corresponds to different tissue types, and the color signifies the storage year. **D.** PCA plots visualizing variance between samples based on tissue types and histologic diagnosis. The accompanying bar chart indicates the index of contribution to these clusters, incorporating clinical information of the samples. Bar color represents the significant contribution (adjusted p value were calculated by Bonferroni method).

### Assessment of in-depth Proteome profile quality

To unravel the influence of prolonged storage period on proteomic data, we partitioned a set of 18 paired tumor samples along with their matched NATs. The samples were divided into two distinct TMT sets: ’Set1’ for samples with long-term storage and ’Set2’ for those with short-term storage, based on the storage duration.

We conducted a comparative analysis by examining the highest-ranking peptides in the global, phosphorylated, and acetylated proteome datasets, based on their PSM counts (**Figure 3A, Supplementary Table 3A**). We focused on the top 1% of features from the overall global and phosphorylated proteome data, while considering the top 300 features from the acetylated proteome dataset for further analysis. The linear regression analysis comparison between the two sets showed R values of 0.75, 0.79, and 0.62 (p < 2.2 X 10^-16^), respectively, indicating that there is no difference between the many observed PSMs produced by the two sets. Nonetheless, a general declining pattern is evident in the ranking of long-term samples (Set2), with a few outliers identified. Specifically, 40 outliers were noted in the global proteome, and 14 outliers were identified in both the phosphor- and acetyl proteomes. Remarkably, the protein P68871 (HBB) emerged as an outlier present in both the global and acetyl datasets. Additionally, we compared the results of PSMs on PTMs with previous studies summarized the PTM in which the most mass shift is found in protein analysis using TMT on CPTAC QC samples^22^. We found that the frequency of modifications slightly increased over the storage period to 1% of the total number of PSMs identified (**Figure 3B, Supplementary Table 3B**). Sodium cation modifications of aspartic (D) and glutamic (E) acids were also identified. However, the frequency of identification of both phosphorylation and acetylation was not affected by the storage period.

**Figure 3.**
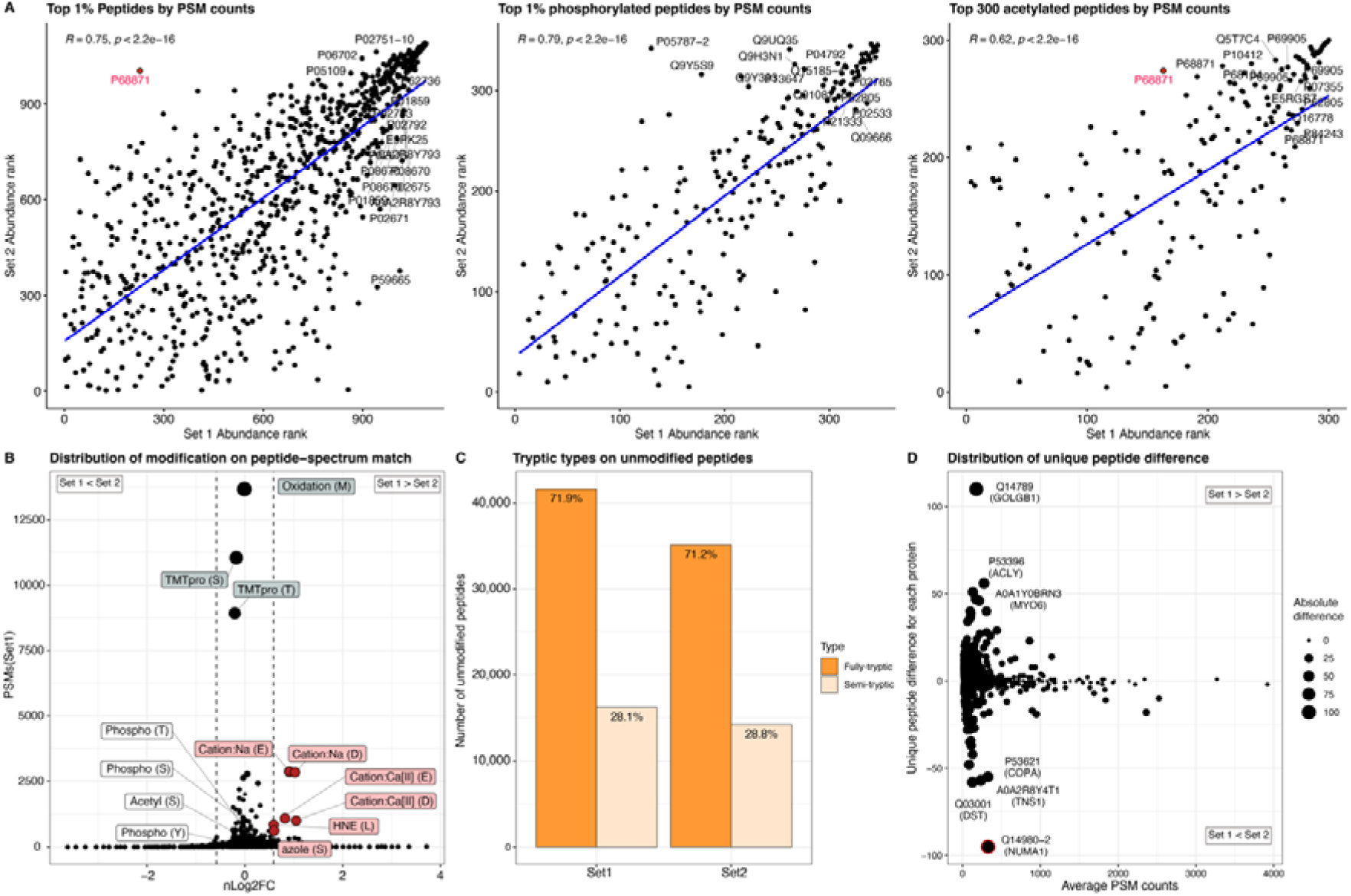
Assessment of proteomic data quality on PSM, peptide, and protein. **A.** Scatterplot illustrating linear regression analysis utilizing peptide and PSM counts from the global proteome and PTM proteome datasets for each TMT set. The regression line is depicted in blue, and the correlation coefficient (R) value is indicated in the upper left corner. Labels denote significant differences based on a rank of 0.1%. **B.** Distribution plot showing identified PTMs in PSMs. The x-axis (nLog2FC) was computed by dividing the PSM counts from Set 1 by those from Set 2 and subsequently normalizing the result using the total PSM count. Significantly differing PTMs are highlighted in red. **C.** Bar chart representing the proportion of tryptic peptide types within unmodified peptides for each TMT set. The color of each bar corresponds to the specific type of tryptic peptide. **D.** Distribution plot depicting the disparity in unique peptide counts between the two TMT sets. The size of each point corresponds to its absolute variation.

Also, we explored the composition of fully tryptic and semi-tryptic peptides in unmodified peptides, as the distribution of tryptic cleaved peptide conformations can change due to storage-induced protein denaturation. Although there was variation in the total number of identified peptides within each set, the relative proportions of tryptic peptides remained consistent across both sets (**Figure 3C**). Fully tryptic peptides assert their dominance, constituting 71.9% and 71.2% within their respective sets. In the context of proteins, the abundance of unique peptides plays a pivotal role in ensuring robustness and accuracy in bottom-up protein qualification and quantification. Among the commonly identified 8,116 proteins, we discover the potential significant discrepancies in unique peptide counts (**Figure 3D**). The proteins which cover low PSM counts have showed a dynamic change in unique peptides. Q14980-2 (NUMA1), which is known as a cancer-associated gene^7^, has a significant low unique peptide in our data. These findings emphasize the feasibility of analyzing tissue samples from different storage periods at various levels of PSMs, peptides, and proteins.

### Expression profile using housekeeping genes and proteins

In tissue sample analysis, the consistent presence of housekeeping genes (HKGs) and proteins (HKPs), as index of cellular equilibrium, offers an essential metric for evaluating sample quality^23^. Their constant and constitutive expression provides a reliable benchmark to assess potential variations resulting from sample collection, preservation, or handling, enhancing the robustness and accuracy. Thus, we collected 802 HKGs from the Housekeeping and Reference Transcript Atlas (HRT Atlas v1.0, www.housekeeping.unicamp.br)^24^ **(Supplementary Table 4A**) and 234 proteins which encompasses 176 ribosomal proteins, 31 RNA polymerase related proteins, and 27 citric acid cycle related proteins from the human protein atlas^25^ **(Supplementary Table 4B**).

We found HKGs were expressed consistently on RNA-seq normalized counts regardless of any clinical information (**Figure 4A**), but HKPs were expressed distinctly across different tissue types (**Figure 4B**). In both cases, the samples predominantly clustered according to tissue type, with the exception of two individual samples: P0008-T for HKG and P0006-T for HKP. The unexpected divergence in housekeeping protein expression between tumor and normal tissue, contrary to the anticipated stable patterns observed in HKGs from RNA-seq data, underscores the intricate dynamics of tumorigenesis^26^. Tumor cells’ unique metabolic demands and uncontrolled proliferation can lead to distinct expression profiles of HKPs involved in energy production and cell cycle regulation. Additionally, the tumor microenvironment and tumor-specific signaling pathways may contribute to the observed variations. These findings reveal a distinct molecular landscape, underscoring the significant role that HKPs play in shaping the unique characteristics of tumor biology. Overall, our analysis indicates that the expression of HKGs and HKPs remains consistent across samples, regardless of the duration of storage, highlighting their potential relevance in elucidating the intricacies of tumor biology through distinct protein expression patterns reflective of tumor characteristics.

**Figure 4.**
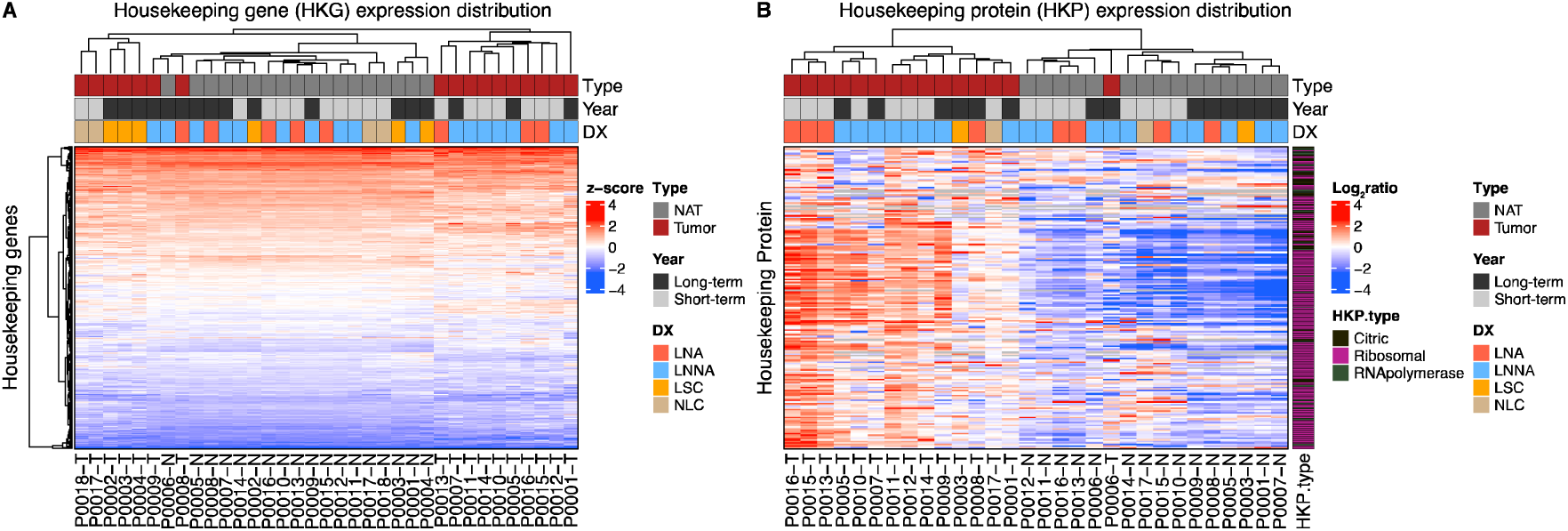
Expression of HKGs and HKPs. **A.** Heatmap depicting the matrix of HKGs using RNA-seq data for each sample. The values in the HKG matrix were presented as z-scores, obtained through transformation from normalized counts using the DESeq2 package in R. **B.** Heatmap illustrating the matrix of HKPs using global proteome data for each sample. In the HKP matrix, values were presented as log_2_ ratios, obtained by calculating the log2 value of each sample’s channel quantification divided by the corresponding CR channel value. Annotations are color-coded to represent tissue type, storage year, histologic diagnosis (DX), and HKP types, including ribosomal proteins, RNA polymerase-related proteins, and citric acid cycle-related proteins.

### Comparison score of hallmark pathways, transcription factors (TFs) and kinases activity

Understanding the complexity of cancer biology is essential for effective therapeutic strategies. Hallmark pathways provide a comprehensive framework to decipher the molecular changes driving tumorigenesis, encompassing processes such as evading growth suppression, and resisting cell death. Also, analyzing TFs and kinases, key regulators of cellular processes, is crucial in unraveling the intricate signaling networks within cells. Dysregulation of these factors can lead to aberrant gene expression and uncontrolled cell growth. Integrating the analysis of hallmark pathways, TFs, and kinases offers a comprehensive view of cancer’s molecular landscape, guiding the development of targeted therapies that address multiple facets of the disease simultaneously, promising improved treatment outcomes.

Based on our prior result (**Figure 2D**), we found that distinctions between normal and tumor tissues surpassed sample storage period, prompting us to focus on inherent tumor tissue disparities. Through a comprehensive exploration, we subjected these tumor tissues to assessment against the hallmark pathways (msigDB.v7.5.1) (**Figure 5A, Supplementary Table 5A**). Our findings predominantly highlighted higher expression in proliferation and signaling groups across tumor samples, underscoring their pivotal role. Moreover, regarding the OXIDATIVE PHOSPHORYLATION within the metabolic group, we observed a distinctive under-expression pattern specific to LNNA in long-term samples. Meanwhile, hallmark pathways related to ’cellular component’, ’development’, ’DNA damage’, ’immune’, and ’pathway’ showed minimal variations, emphasizing their resilience across the tested conditions. This indicates that the majority of the hallmark pathways remain unaffected by the storage period.

**Figure 5.**
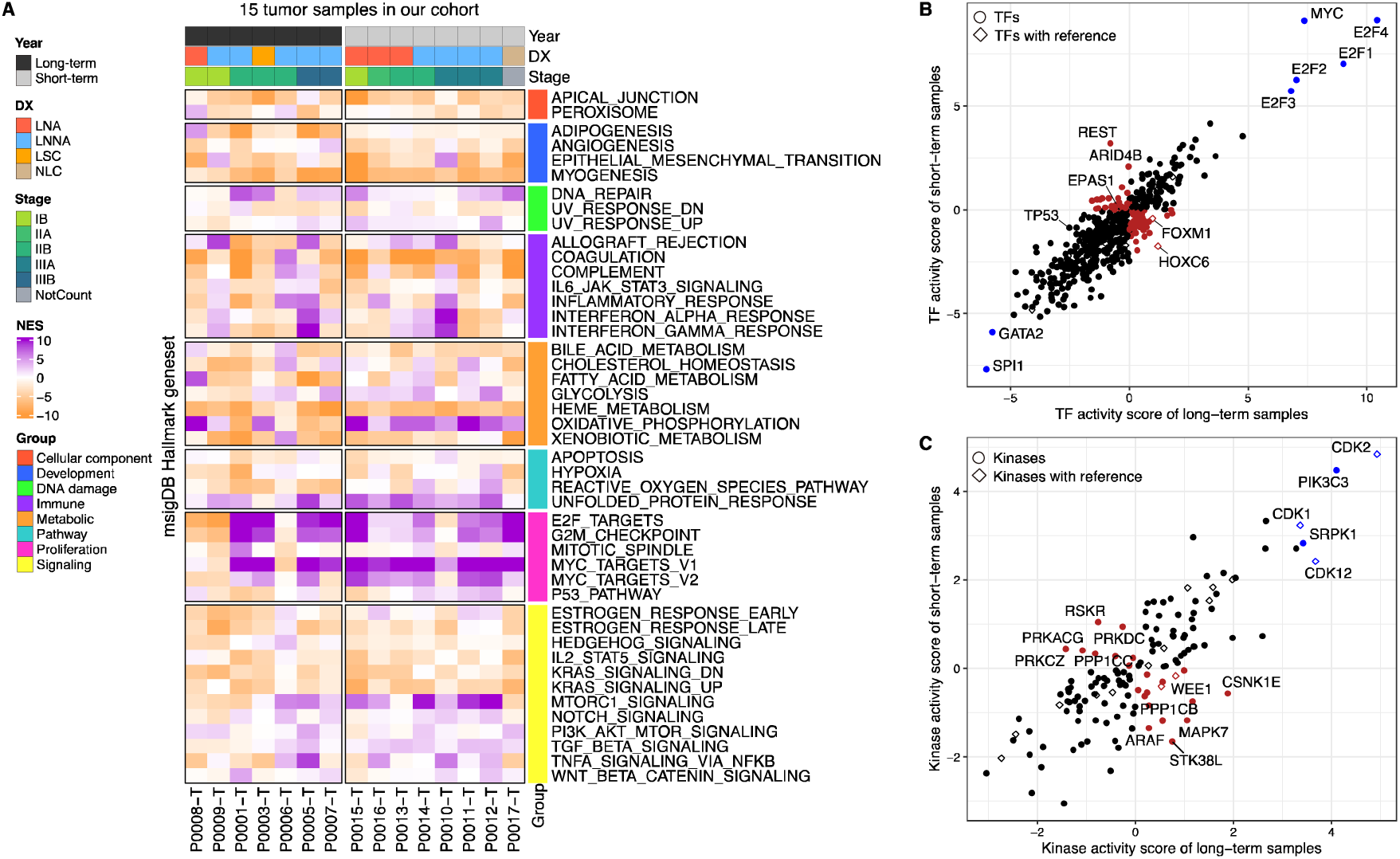
Molecular changes between tumor samples within hallmark pathways, TFs, and kinases activity. **A.** Heatmap illustrating the global protein expression patterns across our tumor samples with respect to msigDB hallmark pathways. Each pathway is grouped and color-coded based on related functions. The Normalized Enrichment Score (NES) is utilized to depict pathway enrichment levels. **B-C.** Comparative analysis of TF and kinase activities between long- and short- term tumors and NAT. In the TF analysis, diamond shapes denote the most frequently dysregulated TFs. In the kinase analysis, diamond shapes represent the list of kinases targeted for inhibition with FDA-approved compounds. The top upregulated and downregulated TFs or kinases are highlighted in blue, while opposite activity regulations in the two sets are indicated by red.

Using differentially expressed genes (DEGs) and phosphosites (DEPPs) expression, we predicted TF activity (**Supplementary Table 5B**) and kinase activity (**Supplementary Table 5C**) by “DoRothEA” and “PHONEMeS” packages in R. A total of 26 TFs associated with lung cancer, encompassing both NSCLC and SCLC^27^, have been identified as frequently dysregulated in lung tumoral cells. Utilizing these TFs, we conducted a comparative analysis between two sample sets, each representing different storage periods (**Figure 5B**). Interestingly, the majority of TFs exhibited consistent alignment regardless of sample storage variation, and the reported TFs displayed moderate activity scores in line with our tissue data. Notably, TFs MYC and the E2F family exhibited elevated activity, reflecting the overall expression patterns related to proliferation and signaling pathways in the tumor tissues examined, in contrast to the NATs (**Figure 5A**). Previously, various kinase inhibitors and their FDA-approved drug targets have been identified^28^. Among these, the CDK family, B-Raf, and SRC were identified in our dataset, showing consistent kinase activity between the two sample sets representing different storage periods (**Figure 5C**). Additionally, the top kinases exhibiting high activity (CDK1, CDK2, PIK3C3) align with global protein expression patterns within the ’PI3K-AKT-MTOR signaling’ and proliferation pathways. Collectively, these findings suggest that comparative analysis of transcriptomic and proteomic data between long-term and short-term samples does not yield discernible variations in the activity of TFs and kinases, which play pivotal roles in modulating downstream pathways essential for shaping the landscape of cancer biology.

## Discussion

As research explores further into the complexities of cancer biology, a potent approach for unraveling the intricate molecular foundations of this intricate disease has surfaced: the integration of multi-omics analyses, which encompass the genome, transcriptome, and proteome. However, the availability of tissue samples poses inherent limitations due to factors such as sample size, collection procedures, and storage duration. Despite these challenges, there is a growing interest in exploring the potential of archived samples to expand the scope of multi-omics research. In this context, our study aims to address the impact of storage period on the genomic, transcriptomic, and proteomic profiles of archived samples, revealing the viability of utilizing such resources for advancing our understanding of cancer biology.

To comprehensively address these trends and limitations, we conducted a systematic examination of archived samples collected over a decade, focusing on their transcriptomic and proteomic attributes. Our analysis revealed complex patterns and dynamic shifts, particularly in the context of transcriptomic data from long-term samples. While a decline in read counts related to protein coding and gene coverage was observed, these changes did not compromise the fundamental gene expression landscape as shown in **Figure 2B-D**. Proteomic inspection encompassed an in-depth exploration of identification results, modification changes, peptide cleavage patterns, and unique peptide counts, all of which collectively indicated that storage period did not exert significant influence on the proteomic profiles as shown in **Figure 3**. Interestingly, comparisons of HKG and HKP expressions yielded consistent transcriptomic levels across samples, while proteomic profiles unveiled distinctive disparities between tumor and normal tissues, potentially indicative of microenvironmental effects as shown in **Figure 4**.

In conclusion, the challenges posed by limited sample availability in multi-omics studies can be partially complemented through the strategic integration of archived samples. While technical shifts were evident in certain aspects of transcriptomic data, the core gene expression patterns remained robust, and the functionality of essential TFs and kinases remained unaffected. These findings emphasize the potential of archived samples as valuable resources for multi-omics research, offering a broader landscape for investigating cancer biology and paving the way for more comprehensive insights into this intricate field.

## Data and Code availability

Proteomic raw data was deposited in K-BDS (Korea BioData Station, https://kbds.re.kr) with the accession ID KAPxxxxx (Now uploading).

Genomic and transcriptomic raw data was deposited in Korean Nucleotide Archive (KoNA, https://kobic.re.kr/kona) with the accession ID, KAPxxxxxx (Now uploading).

No custom code was used or developed for the analyses presented in this study. Standard workflows and open-source R packages and software were used (“Experimental procedures and Statistical Rationale”).

## Supplemental data

This article contains supplemental data.

## Abbreviations

HKG: House keeping gene
HKP: House keeping protein
TF: Transcription factor
SCLC: Small cell lung cancer
NSCLC: Non-small cell lung cancer
ADC: Adenocarcinoma
SC: Squamous cell carcinoma
PSM: Peptide spectrum matched
PTM: Post-translational modification
NAT: Normal adjacent tissues
TMT: Tandem mass tag
CR: Common reference
TMB: Tumor mutational burden
PCA: Principal component analysis
DEG: Differentially expressed gene
DEPP: Differentially expressed phosphosite
NES: Normalized enrichment score

